# Emergence of stealth polymorphs that escape α-synuclein amyloid monitoring, take over and acutely spread in neurons

**DOI:** 10.1101/2020.02.11.943670

**Authors:** Francesca De Giorgi, Florent Laferrière, Federica Zinghirino, Emilie Faggiani, Alons Lends, Mathilde Bertoni, Xuan Yu, Axelle Grélard, Estelle Morvan, Birgit Habenstein, Nathalie Dutheil, Evelyne Doudnikoff, Jonathan Daniel, Stéphane Claverol, Chuan Qin, Antoine Loquet, Erwan Bezard, François Ichas

## Abstract

The conformational strain diversity characterizing α-synuclein (α-syn) amyloid fibrils is possibly at the origin of the different clinical presentations of synucleinopathies. Experimentally, various α-syn fibril polymorphs have been obtained from distinct fibrillization conditions by altering the medium constituents and were selected by amyloid monitoring using the probe Thioflavin T (ThT). We report here that besides classical ThT positive products, fibrillization in saline simultaneously gives rise to competing fibril polymorphs that are invisible to ThT (stealth polymorphs), and that can take over. Due to competition, spontaneous generation of such stealth polymorphs bears on the apparent fibrillization kinetics and on the final plateau values. Their emergence has thus been ignored so far or mistaken for fibrillization inhibitions/failures. Compared to their ThT-positive counterparts, and as judged from their chemical shift resonances fingerprint, these new stealth polymorphs present a yet undescribed atomic organization and show an exacerbated propensity (approx. 20-fold) towards self-replication in cortical neurons. They also trigger a long distance synucleinopathic spread along nigro-striatal projections *in vivo*. In order to rapidly screen fibrillization products for the presence of such stealth polymorphs, we designed a simple multiplexed assay that can be easily and rapidly operated. This assay allows us to demonstrate the sustainability of the conformational replication of these novel and particularly invasive strains. It should also be of help to avoid erroneous upstream interpretations of fibrillization rates based on sole ThT, and to expedite further structural and functional characterization of stealth amyloid assemblies.

**One Sentence Summary:** stealth α-synuclein fibrils take over

---

α-syn is a small synaptic protein normally associated with neurosecretory vesicles as α-helical multimers (*1*) and interacting with VAMP2 (*2*). Under this conformation, α-syn regulates the kinetics of neurotransmitter release in the synaptic cleft (*1,2*). However, before getting associated with the membrane of presynaptic vesicles, α-syn transits in the neuronal cytosol under its free soluble monomeric form (*1,3*). In this latter context, α-syn oscillates between a variety of different conformational states: it is an intrinsically disordered protein (*4*). Certain of these conformations that are enriched in beta folds (*5*) can incidentally get favored and stabilized leading to their self-assembly into growing amyloid structures that progressively attract and wipe out the α-syn produced by neurons. This leads to the formation of long intra-neuronal amyloid fibrils that partly get packed and stored in inclusion bodies called Lewy Bodies (LBs) (*6*), and partly get passed on the neighboring neurons where they propagate the amyloid buildup process (*7,8*), and so on. This elemental self-regenerative process provides an explanation for the progressive intracerebral spread of LBs observed during Parkinson’s disease progression (*9*) and for the gradual worsening of other synucleinopathies (*10*).

Strikingly, during amyloid stacking α-syn can adopt several distinct structural folds resulting in the emergence of fibril polymorphs (*11-17*). Together with their specific fold, their supramolecular properties also get inherited and propagated as the polymorphs self-replicate by templated growth (*18-21*). It is thus tempting to speculate that the distinctive properties of these different polymorphs are causal and bear on the semiology of the different neurodegenerative diseases that are underpinned by a synucleinopathy (*22-24*). We noticed, however, that the biological characterization of structurally defined α-syn polymorphs had often been limited to the demonstration of their cellular “toxicity” or “pro-aggregative” action without putting under scrutiny their ability to reproduce their distinctive amyloid fold inside cells. The original aim of our study was thus to determine whether structurally characterized α-syn polymorphs could demonstrate autonomous self-replication of their structural traits during propagation not only *in vitro* but also in living neurons. During our experiments, we incidentally discovered the existence of stealth polymorphs.

We first sought (i) to isolate/characterize different α-syn polymorphs self-assembled under simple shaking in saline and (ii) to determine if these “untouched” assemblies could self-replicate and spread in neurons (untouched meaning here “without artificially disrupting the assemblies using sonication”). A single batch of « NMR-grade » recombinant α-syn (labeled with ^13^C and ^15^N) *(Methods in Supplementary Materials)* was divided into 7 identical aliquots (Fig. 1A). One was left at 4°C; the other 6 were then shaken together. Strikingly enough, using the classical amyloid probe Thioflavin T (ThT) (*25*) to monitor the fibrillization (*16,17*), we observed a major variability in the development of the ThT signal from one tube to another (Fig. 1B). Certain aliquots showed a very rapid and strong rise in ThT fluorescence, others had a slower increase trend but aiming at comparable final plateau values, while others had an almost flat ThT curve. As it is widely assumed that the buildup of a ThT signal in these conditions is synonymous with proper α-syn amyloid assembly (*16,17*) – see current best practice guidelines – we simply sought to compare the bioactivity of iso1, a presumably “fruitful” ThT-high sample, with that of iso3, a presumably “failed”, ThT-low sample. To that end, we developed and validated a High Content Analysis (HCA) assay based on primary mouse cortical neurons (*27,28*) allowing to observe a seeded synucleinopathic buildup in fully mature primary cultures (30 days *in vitro*) (Fig. S1). The neurons were challenged with iso1 and iso3, immediately upon completion of the 40h fibrillization period without any other manipulation (“untouched”). The results we obtained (Fig. 1C) were completely opposite to our expectations: iso3, which according to ThT was supposed to be virtually devoid of amyloids (*26*), induced a massive synucleinopathy causing a widespread buildup of secondary aggregates, while no synucleinopathy at all developed in the neurons treated with the ThT-positive iso1. To explain this counterintuitive lack of activity, we reasoned that unspecific clumping could have occurred in iso1. Sonication of iso1, which maximally reduced the size of the assemblies (Fig. 1D), did not however overcome the bioactivity deficit of iso1 compared to iso3 (Fig. 1E).

**Fig. 1:**
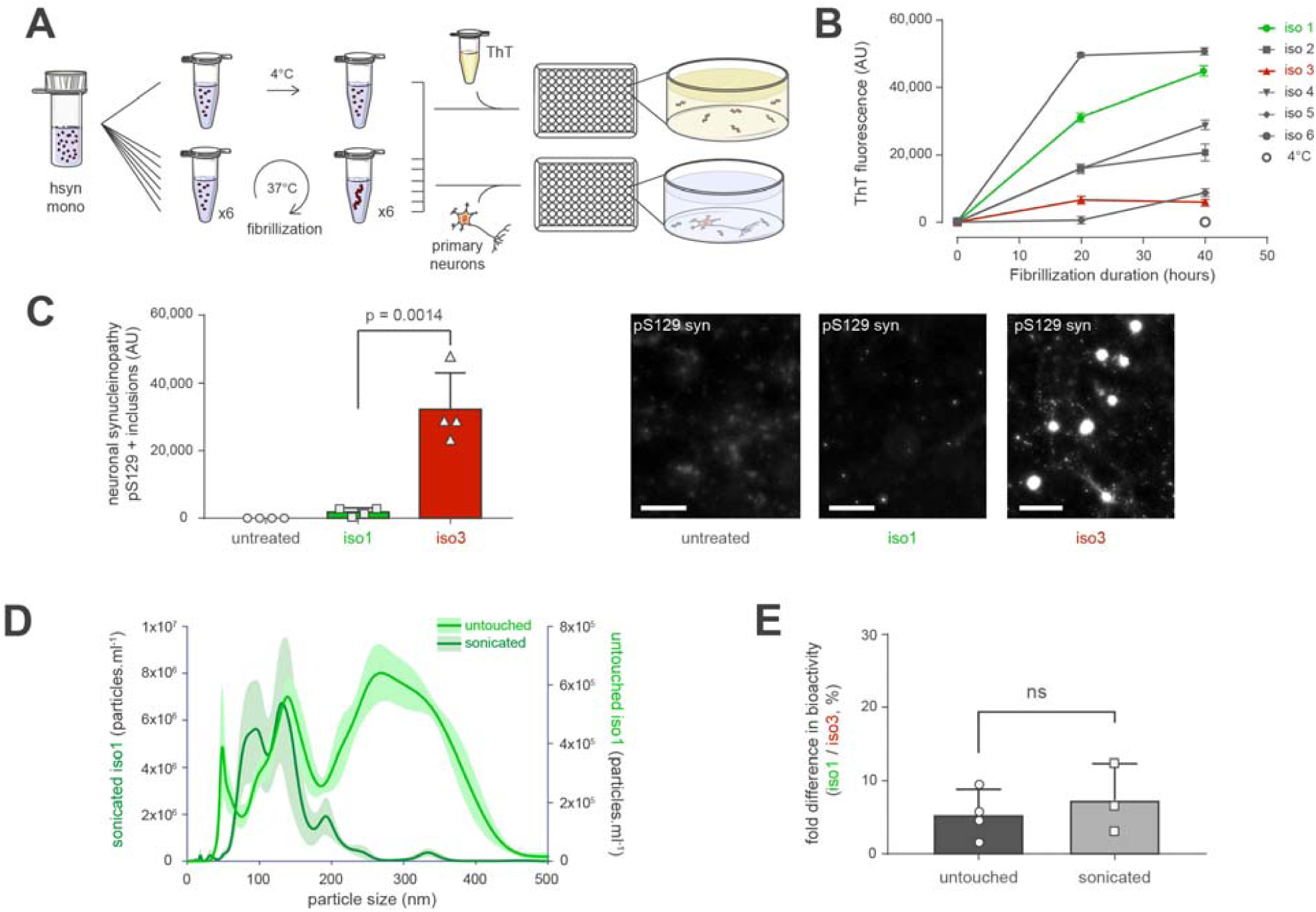
Spontaneous emergence of competing α-syn assemblies that escape amyloid monitoring and show exacerbated spread in neurons. (A) Schematic representation of the fibrillization and testing workflow. (B) Development of the ThT signal in the 6 α-syn tubes, sampled after 0, 20 and 40 hours of shaking at 37°C, or at 40h for the unshaken, 4°C tube. (C) High Content Analysis (HCA) of the synucleinopathy induced by 10 nM (equivalent monomeric α-syn concentration) of untouched iso1 and iso3 in primary mouse cortical neurons; quantitation and representative imaging fields. Scale bar: 100 µm. (D) Nanoparticle Tracking Analysis shows that sonication decreases the size of the assemblies present in iso1. (E) The activity ratio of iso1 to iso3 in the neuronal synucleinopathy assay is insensitive to prior sonication of iso1 and iso3. P values of the group differences calculated using T test.

We thus proceeded with a further comparative characterization of iso1 and iso3 using the Congo-Red derivative X-34 (*29*) (Fig. 2A), density floatation (Fig. 2B), velocity sedimentation (Fig. 2C) (*30*), and eletron microscopy (Fig. 2D). All these approaches indicated that iso3 contained as much of α-syn amyloid assemblies as iso1, with the iso3 fibrils appearing slightly shorter, more straight and bundled at an ultrastructural level (Fig. 2D). Further, while the sedimentation patterns indicated a distinctive size distribution of the assemblies populating iso1 and iso3 (Fig. 2C), the similitude of their density floatation patterns (Fig. 2B) indicated that all the assemblies present in the two preparations shared a common and unique amyloid compactness, irrespective of their particle size. Overall, these data pointed to the possibility that with iso1 and iso3, we were facing two distinct competing amyloid α-syn polymorphs that had spontaneously emerged from a single condition, and endowed with (i) distinct “all-or-none” ThT visibilities, and (ii) distinct “all-or-none” natural propensities to spread in mouse neurons.

**Fig. 2:**
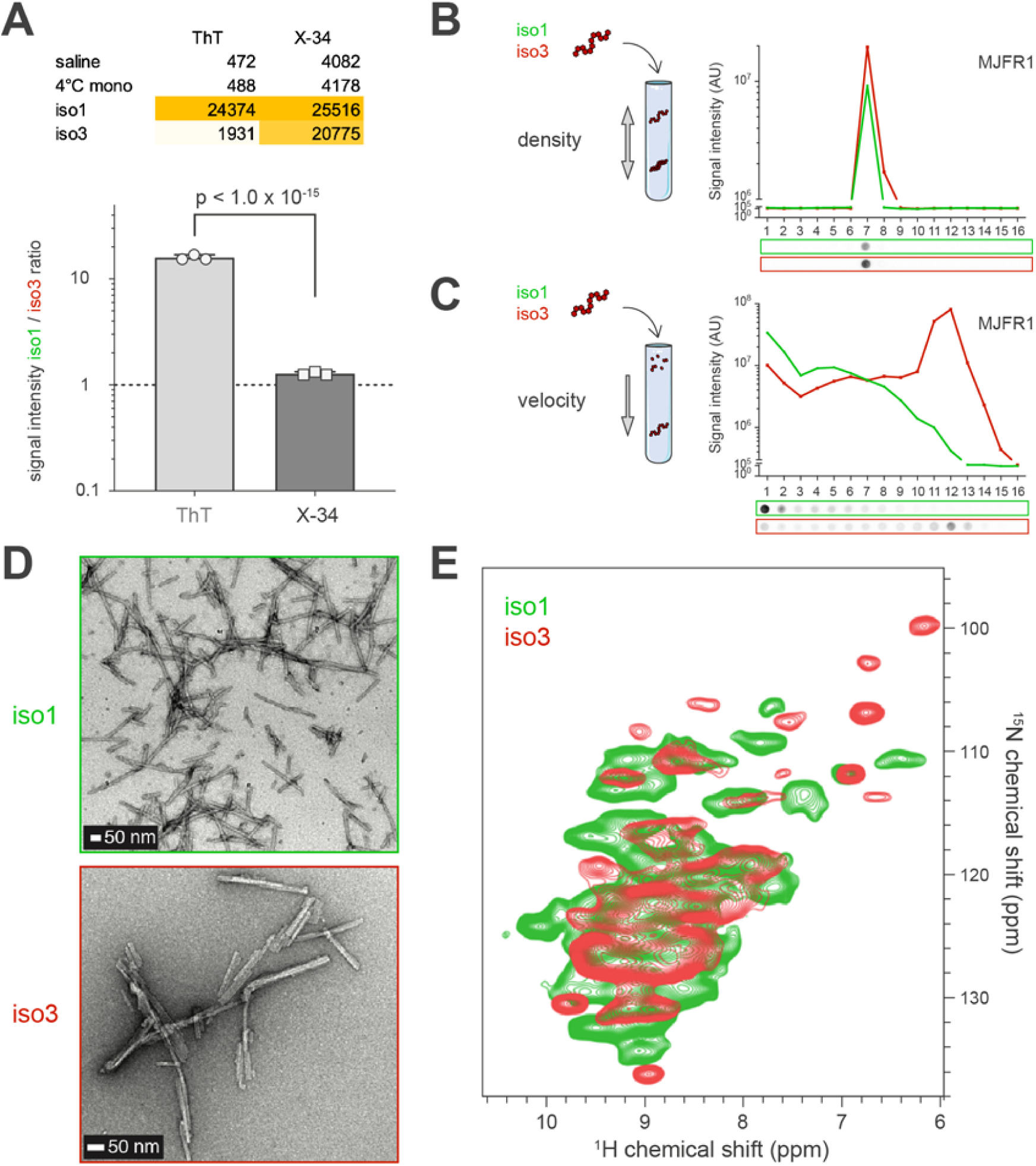
iso3 is a novel stealth fibrillary polymorph. (A) Fluorescence of the amyloid probe X-34 indicates that there is the same amount of amyloids in iso1 and iso3 in spite of an over 10 fold difference in ThT signal; table with mean values (n=3) and corresponding graph with iso1 to iso3 signal ratios. P values of the group differences calculated using T test. (B) Schematic representation of the density floatation protocol: iso1 and iso3 were solubilized with sarkosyl and loaded in the middle of iodixanol gradients and fractionated by density upon isopycnic equilibrium ultracentrifugation. The collected fractions (numbered from top to bottom of the gradient) were analyzed for human α-syn content (using the anti-human α-syn antibody MJFR1) by filter trap (representative pictures of the immunoblots under the graph) by quantification from n=3 replicates, of the absolute chemiluminescence signal (A.U., graph). Data presented are the mean curves (bold lines), with standard deviations (lighter shade areas) for each group. (C) Schematic representation of the sedimentation velocity protocol: iso1 and iso3 were solubilized with sarkosyl and loaded on top of linear iodixanol gradients and fractionated by sedimentation velocity upon ultracentrifugation. The collected fractions were collected and revealed/quantified as in (B). While the density of the iso1 and iso3 assemblies are strictly equal and monodisperse (B), their size distributions are different (C). (D) Transmission electron micrographs of iso1 and iso3 showing their prototypical fibrillary structure. (E) Solid-state NMR hNH experiments of fully protonated iso1 and iso3 showing that they correspond to two distinct amyloid polymorphs at a local level.

We thus performed two-dimensional solid-state Nuclear Magnetic Resonance (NMR) spectroscopy in order to assess the hNH (Fig. 2E) and hCH (Fig. S2A) spectral fingerprints of the fully protonated iso1 and iso3 samples at fast magic-angle spinning (100 kHz). Iso1 and iso3 exhibited two clearly distinct structural conformations at the atomic level, as revealed by two different single sets of chemical shift resonances. Both NMR fingerprints exhibited a single set of sharp signals, indicating the presence in each sample of a single non-polymorphic amyloid structure, highly ordered at the atomic level.

Furthermore, we recorded 3D hCANH experiments on iso1 and iso3 to extract two-dimensional NCA planes and compared them to previous solid-state NMR data of four different polymorphs (*12-15*) spanning the entire structure family spectrum defined by Stahlberg and coworkers (*17*) (Fig. S2B).

Strikingly, iso1 turned out to correspond to the “type 2” fibril polymorph species (*13,17*).

As to the stealth polymorph iso3 instead, we found no match with any of the polymorph structure families previously characterized (*12-17*). Yet, in spite of their general structural dissemblance, iso3 shares its “ThT-invisibility” with the “ribbon” amyloids assembled under salt deprivation (*12,18*). Interestingly, a feature of ribbons that stands out compared to all other fibril types is the particular beta structuration of the N-terminus (*15*): this suggests the possible existence of a similar trait in iso3.

These results indicate that α-syn can spontaneously and randomly assemble into distinct competing amyloids in physiological saline, and that previously unnoticed stealth polymorphs can take over and cause exacerbated neuronal spread.

Additionally, with regards to the current methodological guidelines (*26*), these results warn us about the generic upstream use of ThT to monitor α-syn amyloids: this practice can lead to the conclusion that damped or null fluorescence rates of change correspond to “inhibited” or “failed” fibrillizations, while they can in fact reflect the emergence and growth of stealth polymorphs. This can for instance become a significant concern when studying drug candidates aimed at inhibiting fibrillization. Clearly, upstream screening of fibrillization products with ThT also resulted in the artificial orientation of studies towards a restricted subset of α-syn amyloid structures (*e.g.* Ref. 16).

We thus decided to build a structural assay capable of discriminating both stealth and regular polymorphs and to monitor amyloid species generation. The absence of ThT signal in the iso3 fibrils could derive from only subtle changes in the ThT-binding region of the assemblies. Thus, we hypothesized that fluorescent compounds with a slightly different chemical structure but sharing with ThT its trimethine cyanine skeleton (*31, 32*) could detect iso3. Strikingly, SybrGreen, the well-known DNA intercalating dye (Fig. 3A), not only turned out to be indeed capable of sensing the amyloid structure of iso3, but it also appeared to be much less capable of doing so for iso1, giving an almost mirror-image of the ThT readings: iso1 was ThT-high and SybrGreen-low, while iso3 was ThT-low and SybrGreen-high (Fig. 3B). Instead, the fluorescent Congo-Red derivative X-34, which binds to a distinct region in amyloids (*29, 33*), detected iso1 and iso3 with a similar sensitivity (Fig. 3B): X-34 provided a polymorphism-independent readout of the total amounts of amyloids in each sample. Thus, like for prion strains and LCPs (*34*), multiple external probing using these different molecules could discriminate the α-syn polymorph identities revealed by solid-state NMR. We thus combined these probes into a multiplexed assay to derive a fingerprint of the emerging α-syn assemblies at high throughput and called it the « fibrilloscope » (Fig. 3C), a simple assay with 4 readouts: the 3 dyes ThT, SybrGreen,and X-34; to which we added the measurement of UV Light attenuation to exploit Mie scattering and sense the emergence of particulate assemblies in the 30-300 nm diameter range (*35, 36*).

**Fig. 3:**
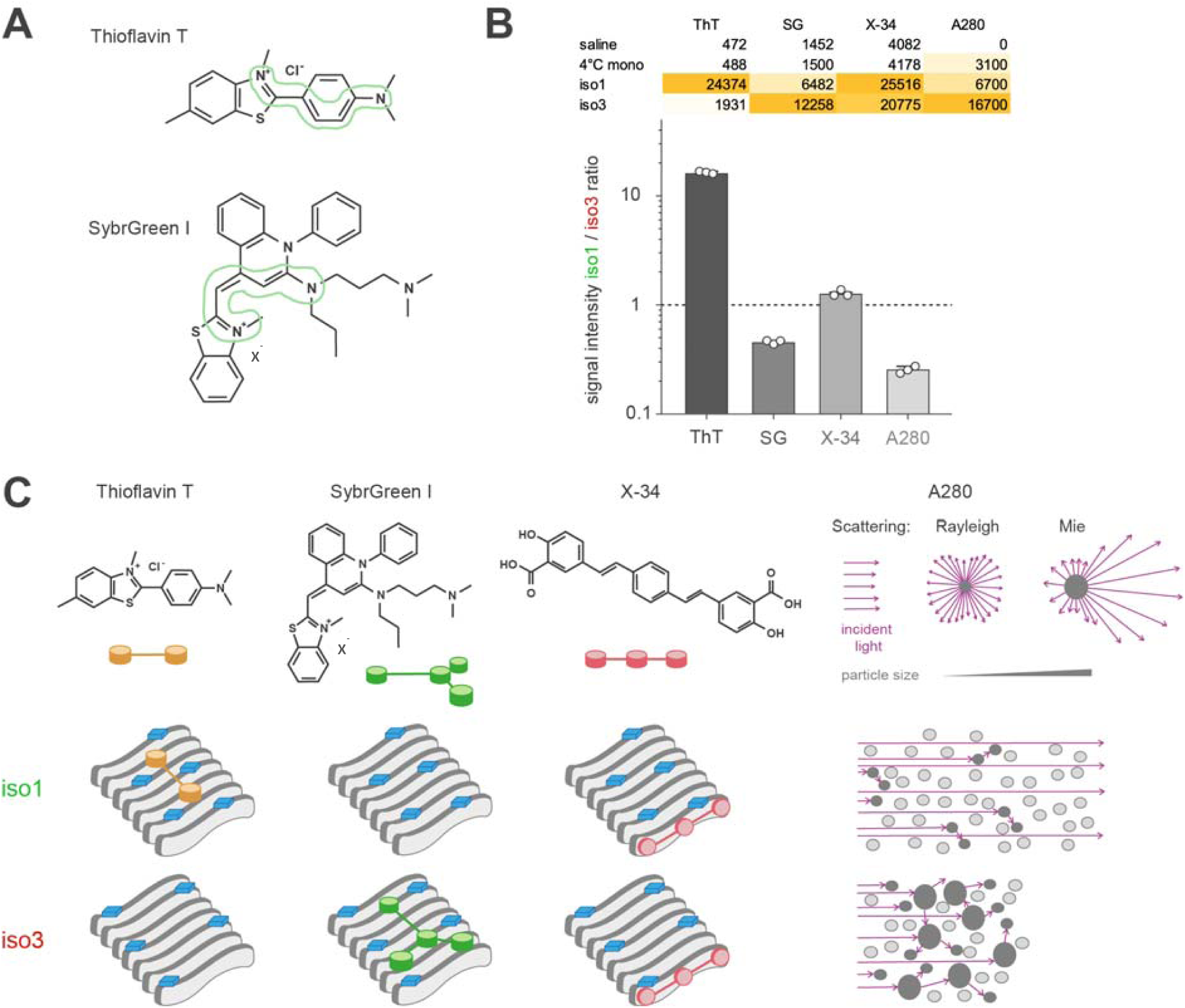
A “fibrilloscope” for rapidly resolving stealth polymorphs. (A) 2D structures of Thioflavin T and SybrGreen I molecules showing their trimethine cyanine skeleton circled in green. (B) Fluorescence intensities of Thioflavin T (ThT), SybrGreen I (SG) and X-34, as well light attenuation at 280 nm (A280) of iso1 versus iso3 samples (A.U., table). Ratios of iso1/iso3 signal intensities were calculated and plotted (bar graph) for each sample and probe, showing a specific detection of iso1 by ThT (>5 times higher than for iso3) and a specific detection of iso3 by SybrGreen I (∼2 times higher than for iso1), and a higher light attenuation (almost 5 times), while detection with X-34 was similar for iso1 and iso3. (C) Tentative model of ThT, SG, and X-34 binding to iso1 (upper) and iso3 (lower schemes). Putative identical exposed residues are depicted in blue bricks, witnessing of a distinct quaternary structure of iso1 and iso3 assemblies and allowing or not the binding of the different probes. The A280 light attenuation difference between iso1 and iso3 is explained by the different relative contributions of Rayleigh (small particles < 30 nm) and Mie (larger particles with diameter from 30 to 300 nm) scattering by the assemblies composing iso1 and iso3.

Using the fibrilloscope we compared a second series of spontaneous fibrillizations, with a series of fibrillizations seeded with 1% of untouched iso1 or iso3 (Fig. 4A). In this second round, we used α-syn monomers bearing a single S129A substitution in order to ascertain that the phosphorylated aggregates appearing during the subsequent biological assays would correspond to assemblies formed at the expense of the endogenous pool of α-syn (*27*) (Fig. S1). Both iso1 and iso3 were able to prompt the appearance and the buildup of amyloids *in vitro* in comparison with all the spontaneous fibrillizations, iso1 being even more potent than iso3 (see the X-34 and UV Light attenuation readouts) (Fig. 4B). Moreover, all the amyloids that developed in the presence of 1% of iso1 (iso1.1 to 1.5) were ThT-high and SybrGreen-low, while all the amyloids observed in the tubes inoculated with 1% of iso3 (iso3.1 to 3.5) were ThT-low and SybrGreen-high (Fig. 4B), corresponding to an exact reproduction of the parental fibrilloscope fingerprint (Fig. 3B). iso1 and iso3 samples thus contained distinctive amyloid particles capable of imposing their own core structural arrangement on monomers by conformational templating, driving the assembly of daughter amyloid structures. Strikingly, the second-generation samples retained the exact biological activity of their parents, i.e., all the samples descending from iso1 were inactive on cortical neurons, and all the iso3 offspring triggered an extensive synucleinopathy (Fig. 4C). This indicated that second to inoculation, amyloid particles proliferated, and that they retained both the fold and the neuronal spread propensity of the parent assemblies.

**Fig. 4:**
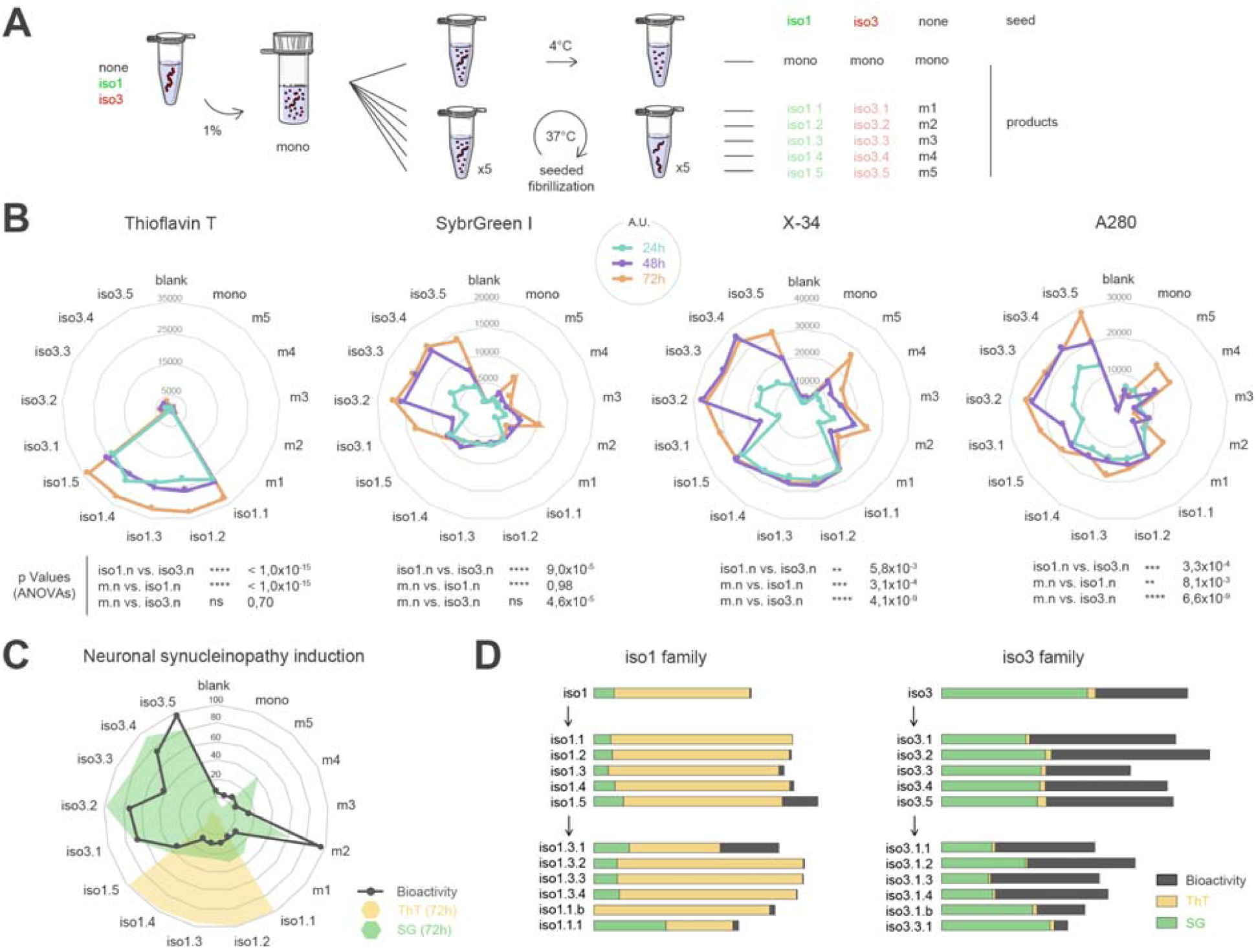
The stealth polymorph iso3 is the founder of a new strain. (A) Schematic representation of the secondary fibrillizations performed in parallel, with and without inoculation, and corresponding nomenclature. (B) Radar plots showing for each individual tube the intensity values of ThT, SG, X-34, and A280 of samples withdrawn at 24h, 48h and 72h during the fibrillization process. The P value of the difference observed with each probe for the different fibrillization groups (1% iso1; 1% iso3; none; n=5 for each group) was calculated using two-way ANOVA followed by Tukey correction and appears under each radar plot. Distinctive kinetics indicative of seeded aggregation and speciation can be appreciated. (C) Radar plot of the extent of the synucleinopathic spread sparked by the 72h secondary fibrillization products in primary cultures of mouse cortical neurons. The latter HCA neuronal bioactivity tests were all run simultaneously, and each individual value plotted in the radar corresponds to the mean of 27 neuronal field quantifications made in 3 replicate culture wells. The corresponding 72h ThT and SG values already shown in panel (B) are superimposed for facilitating the appreciation of correlations. (D) Inheritance of specific ThT, SG and Bioactivity traits across seed generations. In this graph, all 3 parameter values were divided for each single sample by its X-34 value to normalize the parameters with respect to the total amyloid load characterizing each sample. The latter normalization was not explicited in the title blocks for the sake of simplicity. The statistical significance of the inter-generation (children *vs.* grand children for each group) and inter-groups (iso1 *vs.* iso3 for each generation) differences was determined using the Hotelling’s T^2^ test with Bonferroni’s correction for multivariate analysis exploiting simultaneously the 3 parameters characterizing every sample: iso1 children *vs.* iso1 grandchildren: P= 0.70; iso3 children *vs.* iso3 grandchildren: P=0.84; iso1 children *vs.* iso3 children: P<10^−15^; iso1 grandchildren *vs.* iso3 grandchildren: P=0.012.

To explore the sustainability of this particle proliferation we performed a third round of simultaneous fibrillizations in which we compared a series of conditions inoculated with 1% of the second generation fibrillization products (Fig. S3). This resulted in third-generation amyloids that exhibited the same fibrilloscope fingerprint and the same specific bioactivity as their ancestors (Fig. 4E).

These results indicate that the novel stealth polymorph iso3 is capable of replicating its distinctive amyloid fold with the same fidelity and sustainability as iso1, a type 2 polymorph species member (*17*).

We thus looked more closely at the neuronal bioactivity of the different amyloid species that emerged in our experiments. First, we tried to determine why the iso1 strain members, already described as belonging to the type 2 polymorph species (*17*) and which proliferated most efficiently *in vitro*, were unable to spread in neurons. Based on previous observations concerning this polymorph (*22*) we supposed the existence of a monomer concentration threshold to conformational templating and thus forced the mouse neurons to overexpress human α-syn. Strikingly, this overexpression was enough to rescue the bioactivity of iso1 to a level comparable to that of iso3 (Fig. 5A). In addition, comparative mass spectrometry analysis of the neuronal assemblies that formed in these conditions indicated that endogenous mouse α-syn was excluded from the neo-aggregates elicited by iso1.1, while those sparked by iso3.1 contained both human and mouse α-syn (Fig. 5B, C). Thus, beyond a monomer concentration threshold, the inability of the iso1 family members to replicate in mouse neurons was probably also associated with a defective templating of mouse α-syn monomers compared to a more “natural” templating of human α-syn monomers, i.e. this was suggestive of a host-species specificity of the templating process for iso1.

**Fig 5:**
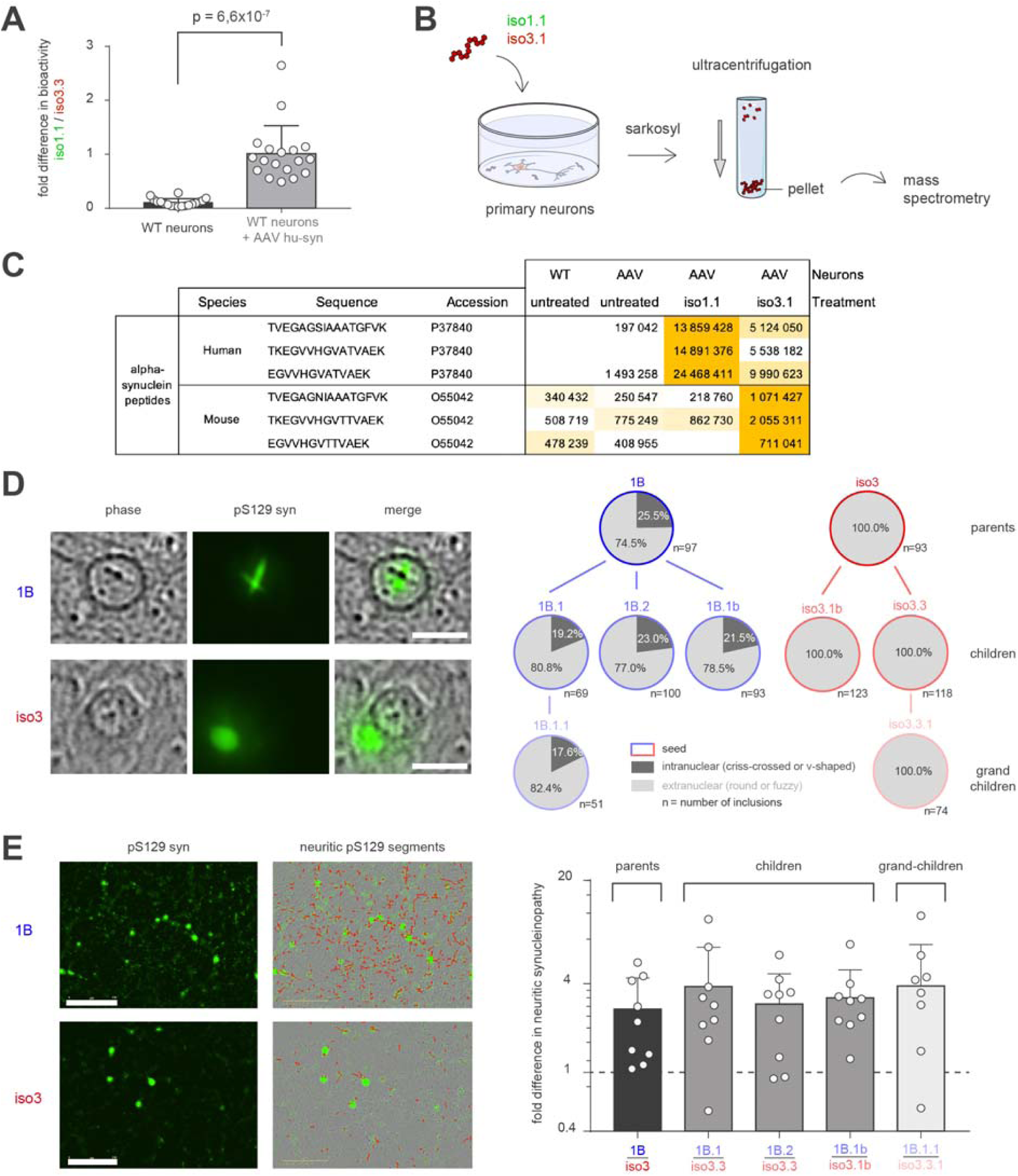
The reference ThT-positive polymorph iso1 needs human α-syn overexpression to demonstrate a bioactivity, while different stealth polymorphs (iso3 & 1B) spontaneously trigger distinct subcellular spread patterns in naïve mouse neurons. (A) Neuronal synucleinopathic spread initiated by iso1 and iso3 was compared in the naïve conditions and in a condition with concurrent infection with an AAV driving the expression of human α-syn by the mouse neurons (see mass spectrometry of panel C). The bioactivity ratio of iso1 to iso3 was dramatically increased in the presence of human α-syn. The P value of the difference in bioactivity observed for the groups (n=18 for each group) was calculated using a T test. (B) Schematic representation of the extraction analysis of insoluble proteins from neurons treated with iso1 and iso3 as in (A). Briefly, mouse primary neurons infected or not with human α-syn AAV and treated or not with iso1 or iso3 were lysed and their insoluble protein content extraction were achieved by solubilization and ultracentrifugation. Pellets containing sarkosyl-insoluble proteins were analyzed by mass spectrometry. (C) Three peptides specific to mouse α-syn, three specific to human α-syn, and five common to both species were identified by mass spectrometry. The species, sequence and hits counts are represented in the table, as well as their sums, represented in the Master protein rows. Mass spectrometry results illustrate the presence of aggregated endogenous mouse α-syn specifically for iso3-treated neurons, while only artificially expressed human α-syn is recovered in the sarkosyl-insoluble fraction from iso1-treated neurons. (D) Left images show typical somatic α-syn inclusions that beacon the experimental synucleinopathy triggered by the stealth polymorphs iso3 and 1B seeds in our HCA assay. Right quantifications show that the intra-nuclear crisscrossed inclusions are distinctive of the synucleinopathy triggered by 1B and its offspring, while they never appear when particles belonging to the iso3 family were used. The counts correspond to the number of somatic aggregates detected in 18 standardized image acquisition fields for each condition. (E) Left images show the extended neuritic synucleinopathy (and its analytical segmentation) caused by 1B compared to iso3 in the neuronal HCA assay. Right quantifications show the 1B/iso3 activity ratio with respect to the induction of neuritic synucleinopathy: this ratio appears to be a constant among all the particle generations produced from 1B and iso3, n=9 for each group, the individual ratio values correspond to the ratio of random pairs of 1B group and iso3 group measurements.

In addition to iso3, we identified a second stealth polymorph 1B also obtained by spontaneous generation in saline. Like iso3, this species was ThT-negative and was able to induce an extensive synucleinopathy in mouse cortical neurons (Fig. S4). However, from a qualitative point of view, the synucleinopathy induced by 1B was characterized by many linear α-syn aggregates in the lumen of the neuronal processes, and by the occasional presence of a single v-shaped or crisscrossed α-syn aggregate embedded in the nuclear chromatin of certain neurons (Fig. 5D, E). In comparison, the synucleinopathy induced by iso3 was characterized by the frequent presence of cytoplasmic round or fuzzy α-syn aggregates located in the soma and of few linear structures in the processes, while intranuclear aggregates were never observed in this case (Fig. 5D, E). The ability to induce these specific patterns was inheritable among seed generations (parents = 1^st^ generation, children = 2^nd^ generation, grandchildren = 3^rd^ generation), and constituted distinctive sustainable traits (Fig. 5D, E). This indicates that like for the type 1 and type 2 polymorph families that have been previously well characterized (*13-17*), stealth polymorphs are also populated by structurally distinct subfamilies, endowed with distinctive and inheritable replication patterns in neurons.

Importantly, the long-range neuronal spread of stealth polymorphs was also observed *in vivo*. Wild-type mice were injected unilaterally at the level of the substantia nigra pars compacta with the 1B polymorph together with an AAV meant to enforce the expression of human α-syn in the nigro-striatal tract. Four months after the unilateral injection, secondary α-syn aggregates were found in the soma of rostral interneurons located in the dorsal striatum, ipsilateral to the injection site (Fig. S5).

These different experiments prompted us to determine whether the aggregates that newly formed in neurons exposed to stealth polymorphs were conformational replicas of the exogenous seeds, in other words, if they resulted from a *bona fide* conformational templating.

To proceed with a comparative analysis of the stealth polymorph iso3 and of its neuronal propagation products we first tried to identify distinctive immunological and physicochemical features that could differentiate iso3 from the regular type 2 polymorph iso1 (*13,17*) and that could be used to characterize the neo-formed assemblies (Fig. 6). In that respect, we observed that the two types of amyloid particles isolated by density floatation exhibited different affinities for the conformation-dependent antibody Syn-F1. The staining of iso1 was much weaker than the one of the stealth polymorph iso3 (Fig. 6A). To determine if this feature was inheritable, we put under scrutiny the daughter amyloid particles iso1.1 and iso3.1 and the grandchild ones iso1.1.1 and iso3.1.1. After simple trapping on nitrocellulose and aldehyde fixation, we found that the two particle species indeed exhibited inheritance of their level of sensitivity towards Syn-F1 (Figs. 6B and S6). We then focused on iso1.1 and iso3.1 and added two upstream steps (i) harsh treatment with sarkosyl mimicking the extraction and recovery procedures of insoluble α-syn amyloids from neurons, and (ii) velocity sedimentation for size separation. These additional steps completely collapsed the Syn-F1 immunoreactivity of iso1.1 while the one of iso3.1 was preserved, further widening the « immunoreactivity gap » between the two species (Fig. 6C).

**Fig 6:**
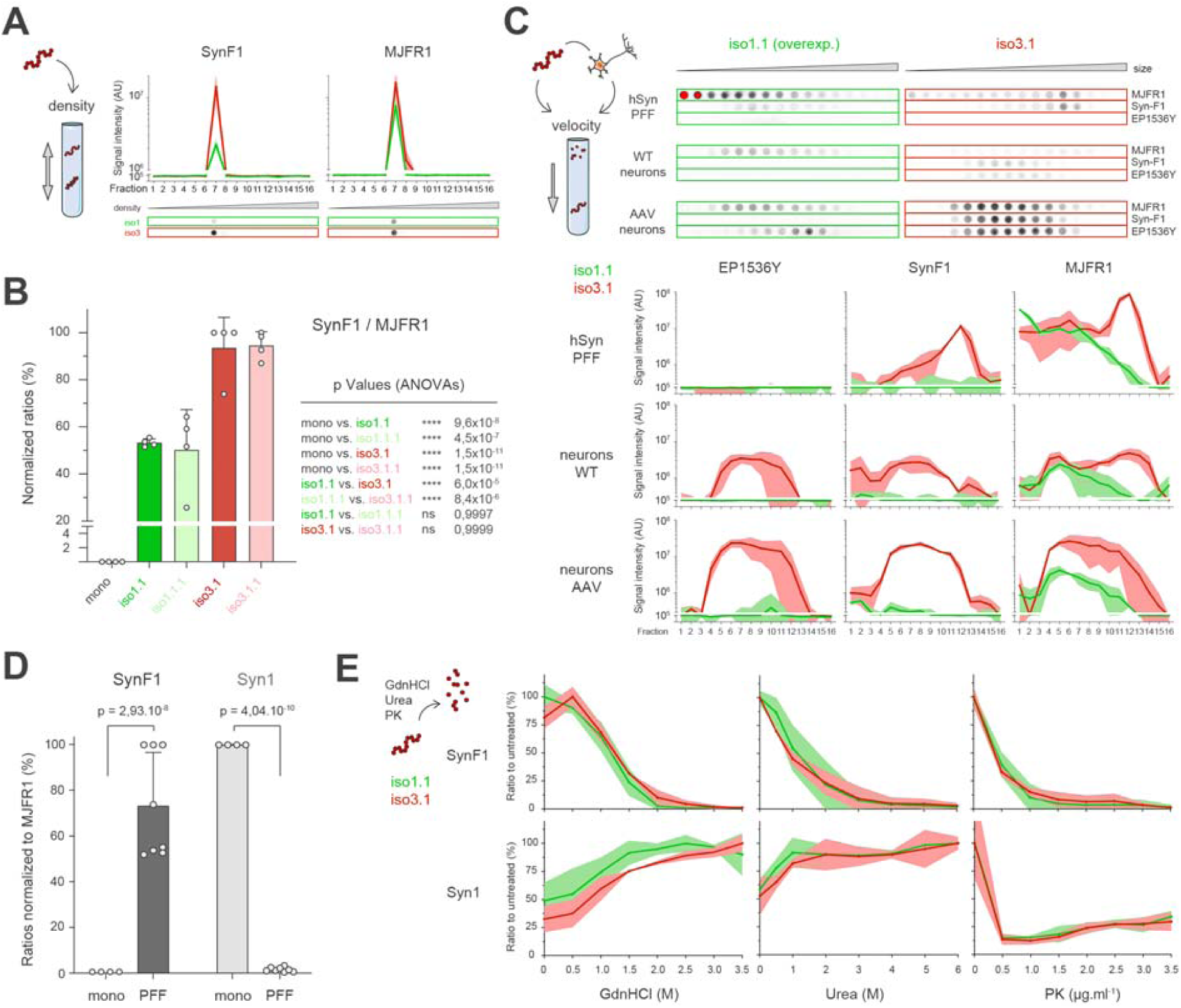
The stealth polymorph iso3 perpetuates its fold in living neurons. (A) Schematic representation of the density floatation procedure applied to iso1 (green) and iso3 (red) preparations. The collected fractions were analyzed for aggregated α-syn (SynF1) and human α-syn (MJFR1) content by filter trap (representative pictures of the immunoblots under the graph) by quantification from n=3 replicates, of the absolute chemiluminescence signal (A.U., graph). Data presented are the mean curves (bold lines), with standard deviations (lighter shade areas) for each group. (B) Monomeric (grey) and iso1.1 (dark green) iso1.1.1 (light green), iso3.1 (dark red) iso3.1.1 (light red) fibrillar recombinant human α-synuclein preparations were subjected to filter trap assays followed by immunoblotting with antibodies against aggregated (SynF1) and total human (MJFR1) α-syn. Ratios of SynF1 to MJFR1 signal intensities are plotted in a bar graph with individual n=4 samples per group. Two-ways ANOVA followed by Tuckey multiple-comparisons tests gave the indicated p values depicting the significant differences of SynF1 immunoreactivity between iso1 and iso3 species. (C) Schematic representation of the sedimentation velocity procedure applied to iso1.1 (green), iso3.1 (red) PFF preparations, and to primary neurons with or without concomitant AAV-mediated overexpression of human α-syn and treated with iso1.1 or iso3.1. The collected fractions were analyzed for human (MJFR1), aggregated (SynF1) and pS129 phosphorylated α-syn by filter trap (representative pictures of the immunoblots, top panel). To note, iso1.1 PFFs and treated neurons sedimentations immunoblots are overexposed compared to iso3.1 in order to allow the visualization of the populations analyzed hereafter. Graphs show the quantification from n=3 replicates per group, of the absolute chemiluminescence signal (A.U.). Data presented are the mean curves (bold lines), with standard deviations (lighter shade areas) for each group. Monomeric and fibrillar recombinant human α-synuclein preparations were subjected to filter trap assays followed by immunoblotting with antibodies to aggregated (SynF1), monomeric (Syn1) and human (MJFR1) α-synuclein antibodies. Ratios of SynF1 and Syn1 to MJFR1 signal intensities are plotted in a bar graph with individual n=4 and n=8 for monomeric and fibrillar groups respectively. Multiple t-tests Holm-Sidak corrected for multiple comparisons gave the indicated p values reflecting the specific immunoreactivity of α-syn fibrils and monomers towards, respectively SynF1 and Syn1. Note that the selectivity of Syn1 towards monomers was previously ignored. (E) Iso1.1 (green) and iso3.1 (red) were subjected to guanidinium (GdnHCl), urea and proteinase K (PK) treatments of increasing concentration (0 to 3.5M GdnHCl for 1 hour at room temperature, 0 to 6M urea for 6 hours at room temperature, and 0 to 3.5µg/ml PK for 30 minutes at 37°C), followed by filter trap assays immublotted with antibodies to aggregated (SynF1, upper graphs) or monomeric (Syn1, lower graphs) α-synuclein. The signal intensity of each point is expressed as a ratio to the untreated sample in the same conditions, allowing to draw curves depicting the disassembly, denaturation and proteolysis of the fibrils, respectively. Data presented are the mean curves (bold lines), with standard deviations (lighter shade areas) for each group of n=3 replicates. Representative filter trap assay immunoblots of each treatment are shown in Fig. S6.

Under such analytical conditions, and as one could have expected at this point, the neo-formed α-syn assemblies that proliferated inside the neurons expressing human α-syn also inherited the specific « all or none » Syn-F1 immunoreactivity of the seed species used to inoculate the cultures (Figs. 6C and S6). This indicated that “seed species”-specific templated growth occurs during propagation inside living neurons, for both the regular type 2 polymorph iso1 (*13,17*) as well as for the stealth polymorphs iso3.

It is worth noting that, apart from its solid-state NMR and fibrilloscope fingerprints and from a distinctive reactivity of its Syn-F1 epitope, the stealth polymorph iso3 was otherwise undistinguishable from iso1 with respect (i) to its sensitivity to proteinase K, (ii) to its resistance to urea or guanidinium chloride (Fig. 6D, E), and (iii) to its gross fibrillary appearance (Fig. 2D). This indicates that both the fibrilloscope fingerprint and the Syn-F1 epitope can bear witness to subtle molecular arrangements not reflected “macroscopically”, yet encoding an amyloid particle size distribution (Figs. 2C & 6C), a particle proliferation rate (Fig. 4B), as well as neuronal and cytopathological spread patterns (Fig. 5) that are distinctive of stealth polymorphs.

Our study demonstrates that spontaneous and uncontrolled emergence of stealth α-syn fibril polymorphs with exacerbated neuronal spreading abilities can “silently” take place during the preparation preformed fibrils (*26*). This is clearly asking the question of whether the bioactivity characterizing such preparations is in fact partly due to the unsuspected presence of stealth polymorphs in variable amounts. This is also prompting to an accelerated investigation of the possible specific role of stealth α-syn polymorphs in synucleinopathies (*37*). Intriguingly, the ThT negativity of α-syn amyloids now appears to repeatedly reconduct to MSA: (i) ribbon amyloids that are ThT-negative (*12, 18*) were proposed to recapitulate signs of MSA in injected mice (*22*), (ii) fibrils amplified from MSA extracts are barely ThT positive compared to those amplified from Parkinson’s disease (*24*), and (iii) we show here that our polymorph 1B (ThT negative) specifically triggers the frequent formation of α-syn inclusions criss-crossing neuronal nuclei. This is reminiscent of the morphology of the Neuronal Nuclear Inclusions that are uniquely seen in MSA and that have been proposed to engage neuronal death (*38,39*). This series of clues certainly point to stealth α-syn amyloid polymorphs as possible specific agents of MSA.

## Acknowledgments

We thank L. A. Largitte, L. Basurco, M. L. Thiolat & T. H. Nguyen for technical help, S. and D. Rowe for revising the English language, and T. Boraud for critical reading of the manuscript.

## Author contributions

conceptualization: FDG, FI; validation, supervision: FDG, EB, FI; investigation, formal analysis: FDG, FL, FI, FZ, ALe, MB, XY, EM, AG, ED, BH, ND, JD, SC, ALo; methodology: FDG, FI, FL, BH, EF, CQ, ALo; visualization: FL, FDG, FI; writing-original draft preparation: FI; writing-review and editing: FDG, FL, EB, BH, ALo, FI.

## Supplementary Materials

### Materials and Methods

#### α-syn expression

*E. coli* strain BL21(DE3) plysS was transformed with pET24-α-Syn vector by electroporation and plated onto LB agar plate containing 30 µg/mL Kanamycin. A pre-culture in 5 mL LB medium was inoculated with one clone and incubated at 37 °C under 200 rpm shaking for 4 hours. The expression on α-syn was carried out in M9 minimal medium containing 2 g/L of ^13^C Glucose and 1 g/L of ^15^NH_4_Cl as carbon and nitrogen sources. Cells from LB pre-culture were recovery by centrifugation (1,000*g – 10 minutes) and used for inoculating 200 mL of M9 medium. Cells were grown overnight at 37 °C under 200 rpm shaking and then diluted in 2 L of culture. Protein expression was induced by adding 1 mM IPTG during exponential phase, evaluated by Optical Density at 600 nm reaching 0.8. Cells were harvested after 4-5 hours of culture at 37 °C by 6,000*g centrifuge (JLA 8.1 Beckman Coulter) and pellet was kept at −20 °C before purification. The site-specific non-phosphorylable mutant S129A was obtained by site-directed mutagenesis of pET24-α-Syn.

#### α-syn purification

Pellet was thaw in 10 mM Tris-HCl (pH 8.0), 1 mM EDTA and 1 mM PMSF, Pierce Complete EDTA-free protease inhibitors tablet (Thermofisher) buffer and sonicated 3 times 45 sec (Bandelin Sonoplus – VS70T probe) previous to be centrifuged. The supernatant was boiled for 20 minutes and centrifuged. Streptomycin sulphate was added to supernatant to final concentration of 10 mg/mL and the solution was stirring for 15 minutes at 4 °C then centrifuge. Ammonium sulphate was added to supernatant to a final concentration of 360 mg/mL and the mixture was stirring to 15 minutes at 4 °C before to be centrifuged. These four centrifugations were performed at 20,000 rpm for 30 minutes and at 4 °C with Beckman Coulter JA-25.5 rotor. The pellet was resuspended in 25 mM Tris-HCl (pH 7.70) and dialysed against the same buffer to eliminate salts. The dialysed sample was injected onto HiTrap Q HP column previously equilibrate with 25 mM Tris-HCl (pH 7.70) and α-synuclein was eluted around 250 µM of NaCl by steps from 0 mM to 500 mM NaCl with AKTA pure system. Fractions containing the protein were dialysed against 20 mM Tris-HCl (pH 7.40), and 100 mM NaCl buffer before to be loading onto HiLoad 26/600 Superdex 75 pg column equilibrate with the same buffer with AKTA pur system. Monomeric fractions were collected and concentrated if needed by using Vivaspin 15R 2 kDa cut off concentrator (Sartorius Stedim). Purification fractions were checked by using Poly Acrylamide Gel Electrophoresis Tris-tricine 13 % dying with ProBlueSafe Strain. Protein concentration was evaluated spectrophotometrically by using absorbance at 280 nm and extinction coefficient of 5,960 M^-1^.cm^-1^.

#### α-syn fibrillization

Solutions of monomeric α-syn at 4-5 mg/ml were sterilized by filtration with 0.22 µm Millipore single use filters and stored in sterile 15 ml conical falcon tubes at 4°C. Sterilized stock was then distributed into safe-lock biopur individually sterile-packaged 1,5 ml Eppendorf tubes as 500 µl aliquots. The tubes were cap-locked and additionally sealed with parafilm. All the previous steps were performed aseptically and in a particle-free environment, under a microbiological safety laminar flow hood. For comparative fibrillizations, all the samples were loaded simultaneously in a Thermomixer (Eppendorf) in a 24 position 1,5 ml Eppendorf tube holder equipped with a heating lid. Temperature was set to 37°C and shaking to 2000 rpm. Sampling for measurements during the fibrillization process was done by temporarily returning the samples under the microbiological safety laminar flow hood.

#### Amyloid probes and Fibrilloscope measurements

Samples from the fibrillized α-syn aliquots or of the control α-syn monomers stored at 4°C were diluted to 0,1 mg/ml in 100µl of PBS containing either 20µM Thioflavin T (ThT), 20µM X-34, 0.1% of SybrGreen I (SG) commercial stock solution, or nothing else. The diluted samples were distributed either in 96 well plates (samples with probes), or in UVettes (Eppendorf) (samples without probe). After 20 min at room temperature protected from ambient light, the plates were read under orbital shaking in a BGM Labtech FLUOstar OPTIMA fluorimeter. Excitation/Emission wavelength pairs (in nm) were: 380/520 for X-34, 450/480 for ThT, and 485/520 for SG. The UVettes were read at 280 nm using the 1 cm optical path with an Eppendorf biophotometer in absorbance mode to generate the A280 light attenuation readout.

#### Sonication and Nanoparticle Tracking Analysis

Samples from the fibrillized α-syn aliquots or of the control α-syn monomers stored at 4°C were diluted to 0.1 mg/ml in 100µl in PBS and distributed in cap-locked sterile 0.5 ml PCR tubes (Thermo Fisher). When relevant, sonication was performed at 25°C in a BioruptorPlus water bath sonicator (Diagenode) equipped with thermostatic control and automated tube carousel rotator. The sonication power was set to “high” and 10 cycles of 30 seconds “on” followed by 10 seconds “off” were applied. The impact of this sonication protocol on the particle contents of the samples was scored using an InSight NS300 Nanoparticle Tracking Analysis platform (Malvern).

#### High Content Analysis of experimental synucleinopathy in neurons

Timed pregnant C57/BL6J female mice were received from Charles River Labs 2 days before initiation of the primary culture. The cultures were synchronized with the α-syn fibrillizations in order to use untouched amyloids in the bioassay. Cortices were harvested from E18 mouse embryos and dissociated enzymatically and mechanically (using the neuronal tissue dissociation kit, C-Tubes, and an Octodissociator with heaters, Miltenyi Biotech, Germany) to yield a homogenous cell suspension. The cells were then plated at 20000 per well in 96-well plates (Corning, Biocoat Poly-D lysine Imaging plates) in neuronal medium (Neuronal Macs medium, Miltenyi Biotech, Germany) containing 0.5% Penicillin/Streptomycin, 0.5 mM alanyl-glutamine, and 2% Neurobrew supplement (Miltenyi Biotech, Germany). The medium was changed by 1/3 every 3 days, until a maximum of 30 DIV. In such cultures, and in control conditions, neurons represented approximately 85-95% of the cell population, thus, in the text, and for the sake of simplicity, they are refered to as “neurons”. After 7 DIV, vehicle, and α-syn amyloids-containing samples were added at a final concentration of 10 nM equivalent monomeric concentration. When relevant, neurons were infected at DIV10 with α-syn AAV particles (MOI 1000). All the antibodies used and shown in the study are summarized in Table S1. All the plates were acquired and analyzed using an Incucyte S3 High Content Imager (Sartorius).

#### Human α-syn AAV particles

Recombinant AAV9-CMVie/SynP-wtsyn-WPRE vector containing the sequence of human α-syn put under control of the human synapsin promoter was produced by polyethylenimine (PEI) mediated triple transfection of low passage HEK-293T /17 cells (ATCC; cat number CRL-11268). The AAV expression plasmid pAAV2-CMVie/hSyn-wtsyn-WPRE-pA was co-transfected with the adeno helper pAd Delta F6 plasmid (Penn Vector Core, cat # PL-F-PVADF6) and AAV Rep Cap pAAV2/9 plasmid (Penn Vector Core, cat # PL-T-PV008). Cells are harvested 72h post transfection, resuspended in lysis buffer (150 mM NaCl, 50 mM Tris-HCl pH 8.5) and lysed by 3 freeze-thaw cycles (37°C/-80°C). The cell lysate is treated with 150units/ml Benzonase (Sigma, St Louis, MO) for 1 hour at 37°C and the crude lysate is clarified by centrifugation. Vectors are purified by iodixanol step gradient centrifugation and concentrated and buffer-exchanged into Lactated Ringer’s solution (Baxter, Deerfield, IL) using vivaspin20 100kDa cut off concentrator (Sartorius Stedim, Goettingen, Germany).The genome-containing particle (gcp) titer was determined by quantitative real-time PCR using the Light Cycler 480 SYBR green master mix (Roche, cat # 04887352001) with primers specific for the AAV2 ITRs (fwd 5′-GGAACCCCTAGTGATGGAGTT-3′; rev 5′-CGGCCTCAGTGAGCGA-3′) on a Light Cycler 480 instrument. Purity assessment of vector stocks was estimated by loading 10 µl of vector stock on 10% SDS acrylamide gels, total proteins were visualized using the Krypton Infrared Protein Stain according to the manufacturer’s instructions (Life Technologies).

#### Preparation of α-syn amyloid assemblies from pure protein samples and from neurons for analytical centrifugations

Stock preparations of amyloid fibrils (4-5mg/ml in TBS) were diluted to 0.1mg/ml final concentration in TBS. The equivalent of the quantity of recombinant human α-syn fibrils used for the treatment of one petri dish of primary neurons (1.66 µg) was diluted in 500 µL final volume with solubilization buffer (SB): 10 mM Tris pH 7.5, 150 mM NaCl, 0.1 mM EDTA, 1 mM DTT, Complete EDTA-free protease inhibitors (Roche) and PhosSTOP phosphatase inhibitors (Roche), with a final concentration of 1% w/v N-lauroyl-sarcosine (sarkosyl, Sigma), 2mM MgCl_2_, and ∼0.5U/µL of Benzonase nuclease (Millipore). For treated primary neurons solubilization, 500 µL of SB buffer was added per petri dish at room temperature. Cells were scraped with a scraper before transferring the lysate in a 1.5ml centrifuge tube. Fibrils or primary neuron lysates were then solubilized by incubating at 37°C under constant shaking at 600 rpm (Thermomixer, Eppendorf) for 45 minutes.

#### Velocity sedimentation and density floatation gradients

Sedimentations velocity and density floatation gradients fractionations were performed as published previously (*26, 28*). Briefly, for velocity sedimentations, a volume of 400 µL was loaded on top of a 11 mL continuous 10-25% iodixanol gradient (Optiprep, 60% w/v iodixanol, Sigma) in SB buffer with 0.5% w/v sarkosyl linearized directly in ultracentrifuge 12 mL tubes (Seton) with a Gradient Master (Biocomp). For density floatation gradients, 400 µL of solubilized material was diluted in SB buffer with 50% iodixanol and 0.5% w/v sarkosyl for making it 40% iodixanol final concentration. This sample-containing cushion was loaded within a 11.4 mL 10-60% discontinuous iodixanol gradient in SB buffer with 0.5% w/v sarkosyl. The gradients were centrifuged at 200,000 g for 2.5 hours at room temperature (velocity) or at 180,000 g for 16 hours at 4°C (density) in a swinging-bucket SW-41 Ti rotor using an Optima LE-80K ultracentrifuge (Beckman Coulter). Gradients were then segregated into 16 equal fractions from the top using a piston fractionator (Biocomp) and a fraction collector (Gilson). Fractions were aliquoted for further analysis of their content by dot blot, immunoblot on SDS-PAGE or native-PAGE. Gradient linearity was verified by refractometry.

#### Analysis of the protein contents of velocity and density fractions by filter trap

For filter trap assays, native fractions were spotted onto nitrocellulose 0.2 µm membranes (Protran, GE) using a dot blot vacuum device (Whatman). Nitrocellulose membranes were fixed for 30 min in PBS with PFA 0.4% v/v (Sigma) final concentration. After three washes with PBS, membranes were blocked with 5% w/v skimmed powder milk in PBS-Tween 0.5% v/v and probed with primary and secondary antibodies in PBS-Tween with 4% w/v BSA (Supplementary Table 1). Immunoreactivity was visualized by chemiluminescence (Biorad). The amount of the respective protein in each fraction was determined by the Image Studio Lite software, after acquisition of chemiluminescent signals with a Chemidoc digital imager (Biorad). The profiles obtained were normalized and plotted with SD, with all respective student test and ANOVA using the Prism software.

#### Extraction of insoluble proteins from PFF treated primary neurons for mass spectrometry

Fibril-treated primary neurons lysates were prepared from a petri dish as described previously for velocity and density gradients. For extraction of insoluble proteins, 500 µL solubilized lysates were mixed 1:1 with SB 40% w/v sucrose, without sarkosyl MgCl_2_ and benzonase, in 1 ml thickwall polycarbonate ultracentrifuge tubes (Beckman) and centrifuged at 250,000 g for 1 hour at room temperature with a TLA 120.2 rotor using an Optima XP benchtop ultracentrifuge (Beckman). Supernatant were collected by pipetting. Pellets were resuspended in 50 µL PBS and total protein concentration was determined by BCA assay (Pierce) prior to equalization and denaturation for 5 min at 100°C in Laemmli buffer.

#### Mass Spectrometry

##### Sample preparation and protein digestion

10 µg of total protein per sample were deposited onto SDS-PAGE gel for concentration and cleaning purpose. Separation was stopped once proteins have entered resolving gel. After colloidal blue staining, bands were cut out from the SDS-PAGE gel and subsequently cut in 1 mm x 1 mm gel pieces. Gel pieces were destained in 25 mM ammonium bicarbonate 50% Acetonitrile (ACN), rinsed twice in ultrapure water and shrunk in ACN for 10 min. After ACN removal, gel pieces were dried at room temperature, covered with the trypsin solution (10 ng/µl in 50 mM NH4HCO3), rehydrated at 4 °C for 10 min, and finally incubated overnight at 37 °C. Spots were then incubated for 15 min in 50 mM NH4HCO3 at room temperature with rotary shaking. The supernatant was collected, and an H2O/ACN/HCOOH (47.5:47.5:5) extraction solution was added onto gel slices for 15 min. The extraction step was repeated twice. Supernatants were pooled and dried in a vacuum centrifuge. Digests were finally solubilized in 0.1% HCOOH.

##### nLC-MS/MS analysis and Label-Free Quantitative Data Analysis

Peptide mixture was analyzed on an Ultimate 3000 nanoLC system (Dionex) coupled to an Electrospray Orbitrap Fusion™ Lumos™ Tribrid™ Mass Spectrometer (Thermo Fisher Scientific). Ten microliters of peptide digests were loaded onto a 300-µm-inner diameter x 5-mm C18 PepMapTM trap column (LC Packings) at a flow rate of 10 µL/min. The peptides were eluted from the trap column onto an analytical 75-mm id x 50-cm C18 Pep-Map column (LC Packings) with a 4–40% linear gradient of solvent B in 105 min (solvent A was 0.1% formic acid and solvent B was 0.1% formic acid in 80% ACN). The separation flow rate was set at 300 nL/min. The mass spectrometer operated in positive ion mode at a 1.8-kV needle voltage. Data were acquired using Xcalibur 4.1 software in a data-dependent mode. MS scans (m/z 375-1500) were recorded at a resolution of R = 120 000 (@ m/z 200) and an AGC target of 4 × 105 ions collected within 50 ms. Dynamic exclusion was set to 60 s and top speed fragmentation in HCD mode was performed over a 3 s cycle. MS/MS scans with a target value of 3 × 103 ions were collected in the ion trap with a maximum fill time of 300 ms. Additionally, only +2 to +7 charged ions were selected for fragmentation. Others settings were as follows: no sheath nor auxiliary gas flow, heated capillary temperature, 275 °C; normalized HCD collision energy of 30% and an isolation width of 1.6 m/z. Monoisotopic precursor selection (MIPS) was set to Peptide and an intensity threshold was set to 5 × 103.

##### Database search and results processing

Data were searched by SEQUEST through Proteome Discoverer 2.3 (Thermo Fisher Scientific) against the Mus musculus Reference Proteome Set (from Uniprot 2019-07; 55121 entries). Spectra from peptides higher than 5000 Da or lower than 350 Da were rejected. The search parameters were as follows: mass accuracy of the monoisotopic peptide precursor and peptide fragments was set to 10 ppm and 0.6 Da respectively. Only b- and y-ions were considered for mass calculation. Oxidation of methionines (+16 Da) and protein N-terminal Acetylation (+42Da) were considered as variable modifications and carbamidomethylation of cysteines (+57 Da) as fixed modification. Two missed trypsin cleavages were allowed. Peptide validation was performed using Percolator algorithm (L Käll, J Canterbury, J Weston, W S Noble and M J MacCoss. Semi-supervised learning for peptide identification from shotgun proteomics datasets, Nature Methods 4:923 – 925, November 2007) and only “high confidence” peptides were retained corresponding to a 1% False Positive Rate at peptide level. Peaks were detected and integrated using the Minora algorithm embedded in Proteome Discoverer. Proteins were quantified based on unique peptides intensities. Normalization was performed based on total protein amount. Protein ratios were calculated as the median of all possible pairwise peptide ratios. A t-test was calculated based on background population of peptides or proteins. Quantitative data were considered for proteins quantified by a minimum of 2 peptides, fold changes above 2 and a statistical p-value lower than 0.05.

#### Measurements of the resistance of fibrils to disassembly, denaturation and proteolysis

Fibrils stock preparations (4-5mg/ml in TBS) were diluted to 0.1 mg/ml final concentration in TBS. For each guanidine hydrochloride (guanidinium, GdnHCl, Sigma), urea (Sigma) and Proteinase K (PK, Sigma) concentration, 10 µL (1 µg) of PFF were mixed 1:1 with 10 µL stock solutions to make GdnHCl 0, 0.5, 1, 1.5, 2, 2.5, 3, 3.5 M final, urea 0, 0.5, 1, 2, 3, 4, 5, 6 M final, and PK 0, 0.5, 1, 1.5, 2, 2.5, 3, 3.5 µg/ml final concentrations. Mixtures were vortexed gently, prior to incubation for 1h at room temperature for GdnHCl, 6h at room temperature for urea, and 30 minutes at 37°C for PK. At the end of the incubation period, treatments were stopped by quickly diluting the samples with 500 µL PBS and directly subjecting them (100 µL per immunoblotting antibody) to filter-trap assay as described previously. The different a-syn species were quantified by immunolabelling with conformation specific, monomeric or pan a-syn antibodies (Supplementary Table 1), and expressed as a percentage of related untreated samples, allowing to draw curves of PFF disassembly, denaturation and proteolysis with GdnHCl, urea and PK respectively.

#### *in vivo* experimental synucleinopathy

WT mice (2 months old) received unilaterally 1 μl of human α-syn AAV virus (concentration: 4.05 × 10^13^ gcp/ml) mixed either with 1 µl of sonicated stealth polymorph α-syn fibrils 1B (5 mg/ml) or with 1µl of saline by stereotactic delivery to the region immediately above the right SN (coordinates from bregma: AP, –2.9, L, –1,3, DV, –4.5) at a flow rate of 0.4 μl/min, and the pipette was left in place for 5 minutes after injection to avoid leakage. Animals were euthanized after 4 months. Ten mice were used in each group — male and female mixed. The brains were perfused with saline, postfixed for 3 days in 10 ml of 4% paraformaldehyde at 4°C, cryoprotected in gradient 20% sucrose in PBS before being frozen by immersion in a cold isopentane bath (–60°C) for at least 5 minutes, and stored immediately at –80°C until sectioning for immunofluorescence analysis. After serial sectioning, the sections were strained using the following primary antibodies: Syn1 (clone 42, BD) for detecting soluble mouse and human α-syn (dilution 1/500), and EP1536Y (Abcam) for detecting phospho S129 positive amyloid aggregates (dilution 1/500). Syn1 was revealed with an anti-mouse Alexa 488 and EP1536Y with and anti-rabbit Alexa 594. The slides were acquired using an Incucyte S3 High Content Imager (Sartorius) with a home-made 3D-printed slide holder.

#### Electron microscopy

Aliquots of fibrillized samples were diluted with water and one drop was applied to glow-discharged 300-mesh carbon-coated copper grids for 1 minute. Grids were washed with one drop of water before being negatively stained with 2 % Uranyl acetate and dried for 5 minutes in the dark. Samples were observed using a Philips CM 120 (high voltage 120 kV – Lanthanum hexaboride filament) transmission electron microscope in low dose mode. Images were acquired using a Gatan US1000 (CCD) camera.

#### Solid-state NMR

All spectra were recorded on Bruker NEO 800 MHz (^1^H Larmor frequency) spectrometer with an ultra-fast Bruker 0.7 mm HCND probe. The 0.7 mm rotor was placed into a 1.3 mm rotor with rubber plug from the bottom. Approximately ∼200 uL of fibrils resuspended in H_2_O was pipetted in a funnel of filling tool and it was spun for ∼18 h at 12 °C at 29 500 rpm (with the SW32Ti Beckmann ultracentrifugation rotor). The detailed NMR experimental parameters are described in the table below.

|  | iso 1 |  | iso 3 |  |
| --- | --- | --- | --- | --- |
| Experiment | 2D <u>hNH</u> | 2D <u>hCH</u> | 2D <u>hNH</u> | 2D <u>hCH</u> |
| MAS frequency/kHz | 100 | 100 | 100 | 100 |
| Field/T | 18.8 | 18.8 | 18.8 | 18.8 |
| Transfer I | HN-CP | HC-CP | HN-CP | HC-CP |
| $^1\text{H}$ field/kHz | 80 | 80 | 80 | 80 |
| $^{15}\text{N}$ field/kHz | 20 | | 20 | |
| $^{13}\text{C}$ field/kHz | | 20 | | 20 |
| Shape | ramp $^1\text{H}$ | ramp $^1\text{H}$ | ramp $^1\text{H}$ | ramp $^1\text{H}$ |
| Time/<u>ms</u> | 1500 | 300 | 900 | 300 |
| Transfer II | NH-CP | CH-CP | NH-CP | CH-CP |
| $^1\text{H}$ field/kHz | 80 | 80 | 80 | 80 |
| $^{15}\text{N}$ field/kHz | 20 | | 20 | |
| $^{13}\text{C}$ field/kHz | | 20 | | 20 |
| Shape | ramp $^1\text{H}$ | ramp $^1\text{H}$ | ramp $^1\text{H}$ | ramp $^1\text{H}$ |
| Time/<u>ms</u> | 500 | 300 | 800 | 300 |
| $^{13}\text{C}$ carrier/ppm | | 54 | | 54 |
| $^1\text{H}$ decoupling | sITPPM12 | sITPPM12 | sITPPM12 | sITPPM12 |
| $^1\text{H}$ decoupling field/kHz | 25 | 25 | 25 | 25 |
| t1 increments | 74 | 300 | 74 | 300 |
| Windows function | QSine3(4) | QSine3(4) | QSine3(4) | QSine3(4) |
| Sweep width (t1)/kHz | 42.4 | 16.1 | 42.4 | 16.1 |
| Acquisition time (t1)/<u>ms</u> | 11.4 | 9.3 | 11.4 | 9.3 |
| $^1\text{H}$ decoupling | WALTZ16 | WALTZ16 | WALTZ16 | WALTZ16 |
| $^1\text{H}$ decoupling field/kHz | 10 | 10 | 10 | 10 |
| t2 increments | 2048 | 2048 | 2048 | 2048 |
| Windows function | <u>QSine 3(4)</u> | <u>QSine 3(4)</u> | <u>QSine 3(4)</u> | <u>QSine 3(4)</u> |
| Sweep width (t2)/kHz | 100 | 37 | 100 | 37 |
| Acquisition time (t2)/<u>ms</u> | 10.2 | 27.6 | 10.2 | 27.6 |
| Water suppression scheme | MISSISSIPPI | MISSISSIPPI | MISSISSIPPI | MISSISSIPPI |
| Inter-scan delay/s | 1.3 | 2 | 1.3 | 2 |
| Number of scans | 64 | 32 | 64 | 32 |
| Measurement time/h | 2 | 6 | 2 | 6 |

**Fig. S1.**
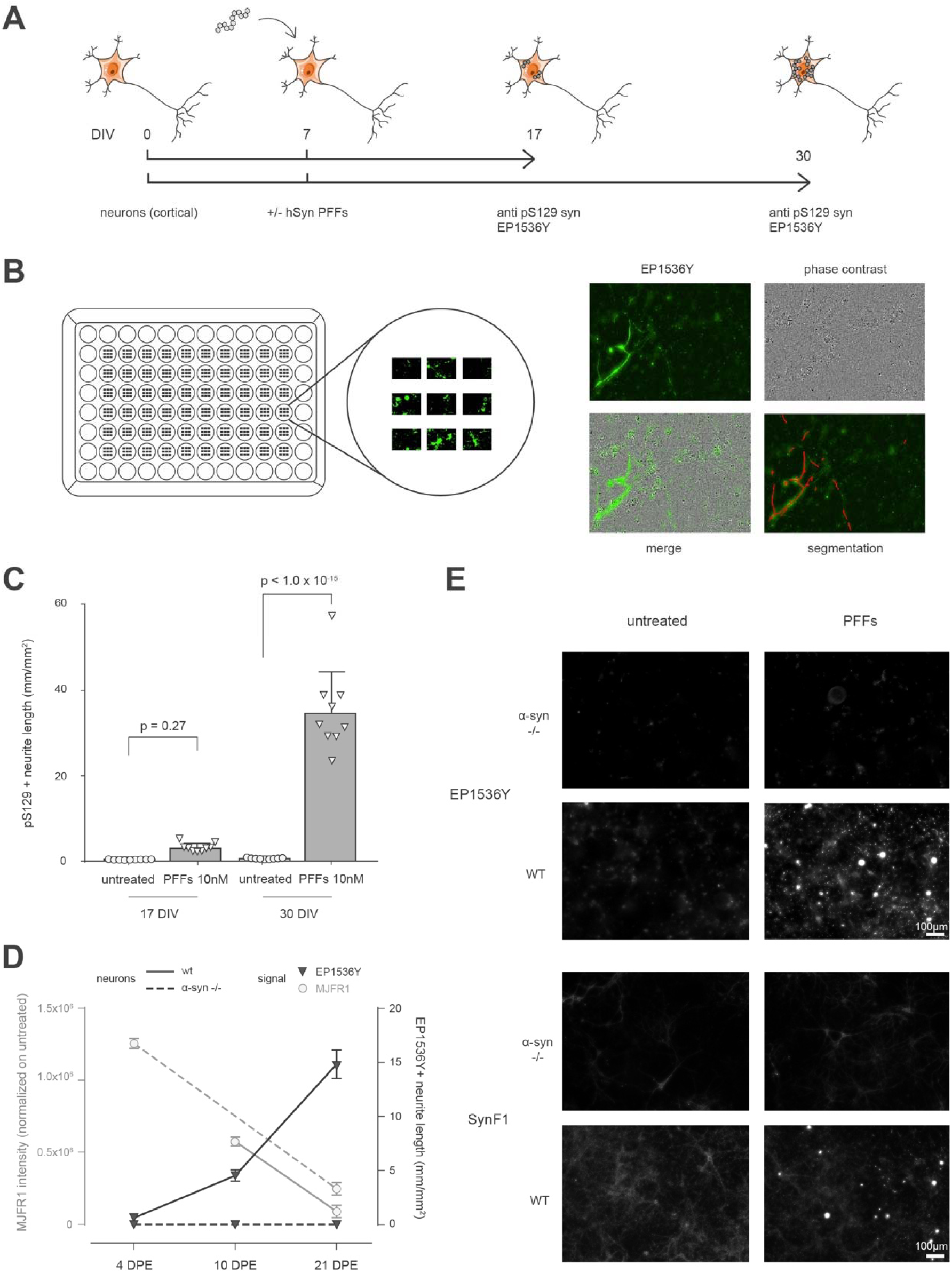
High Content Analysis of experimental synucleinopathy in 96 well primary cultures of mouse cortical neurons. (A) Time schedules of the experiments shown in this figure. (B) Left: schematic view of the seeding plate map and position of the nine 20X imaging fields; Right: representative field of acquisition with single fluorescence and phase contrast channels, corresponding overlay, and overlay with fluorescence segmentation generated for quantitation. (C) Quantification of the experimental synucleinopathy induced by 10 nM α-syn PFFs (batch 1B) equivalent monomer concentration at 17 and 30 days in vitro (DIV) for a common α-syn challenge with PFFs at DIV7. The statistical significance of the comparisons was computed using ANOVA. (D) Monitoring of the progressive degradation of the exogenous α-syn PFFs (MJFR1 positive) and of the concomitant buildup of the neuronal synucleinopathy (EP1536Y positive) in WT and α-syn (-/-) neurons. No synucleinopathic buildup takes place in neurons form α-syn (-/-) mice (Envigo) (DPE: days post PFFs exposure). (E) The synucleinopathy revealed by EP1536Y is also positive for the conformation-dependent amyloid-specific Syn-F1 antibody and corresponds to the recruitment of endogenous α-syn into neo-formed amyloids that get phosphorylated at S129. All the experiments of the paper were analyzed at DIV30.

**Fig. S2.**
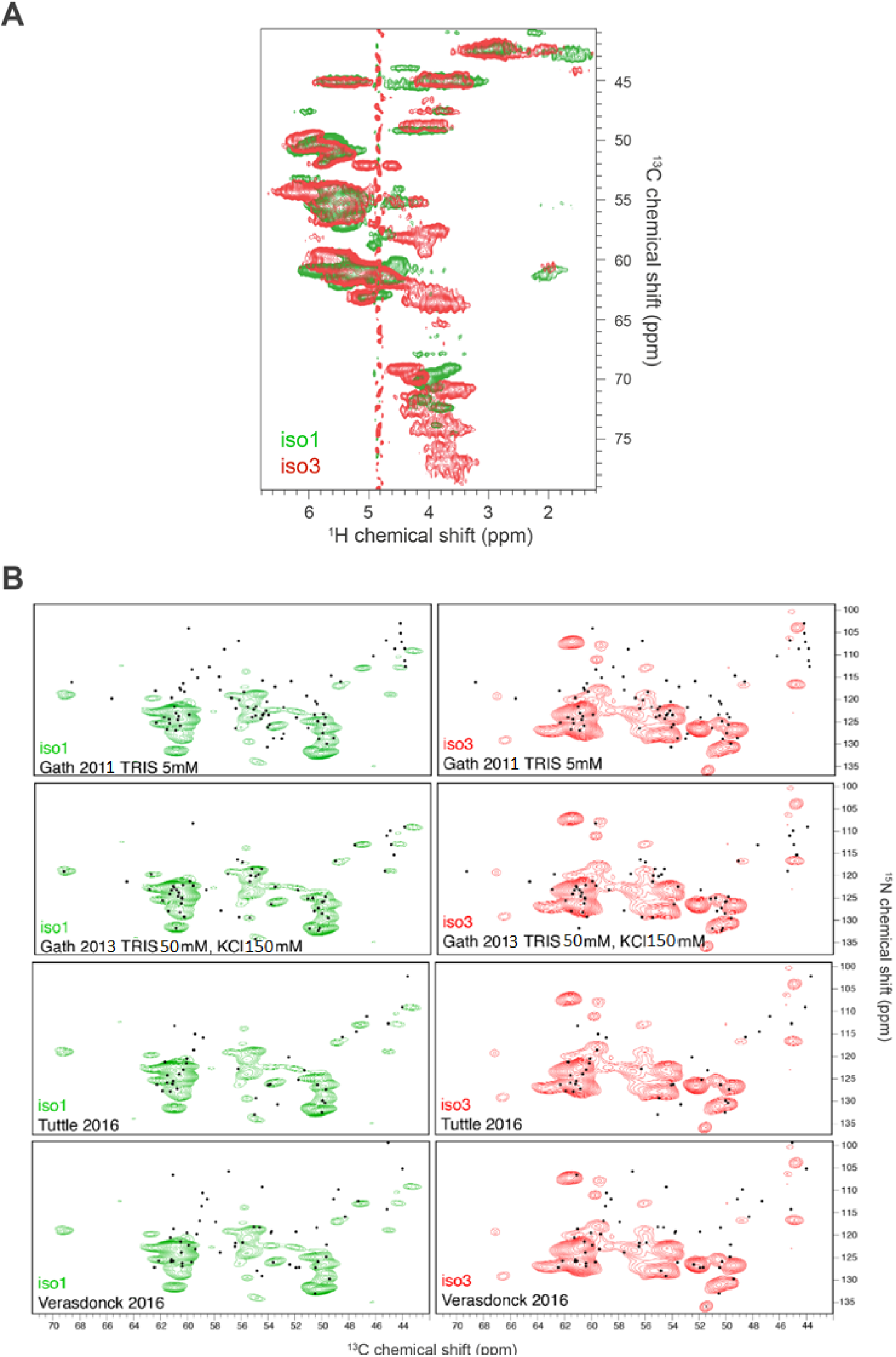
Solid State NMR. **(A)** Solid-state NMR hCH experiments of fully protonated iso1 and iso3 showing that they correspond to two distinct polymorphs. **(B)** Comparison of the 2D NCA projection for the 3D hCANH experiments of iso1 (in green) and iso3 (in red) with reported chemical shift resonances (black dots) of the α-syn fibril polymorphs from references 12 to 15. Note that iso1 is equivalent to the type 2 fibril polymorph (*13, 17*). According to the classification of Ref. 17: Tuttle 2016 and Verasdonck 2016 = Type 1; Gath 2013 = Type 2, Gath 2011 = neither Type 1 nor Type 2 (also known as as “Ribbons”).

**Fig. S3.**
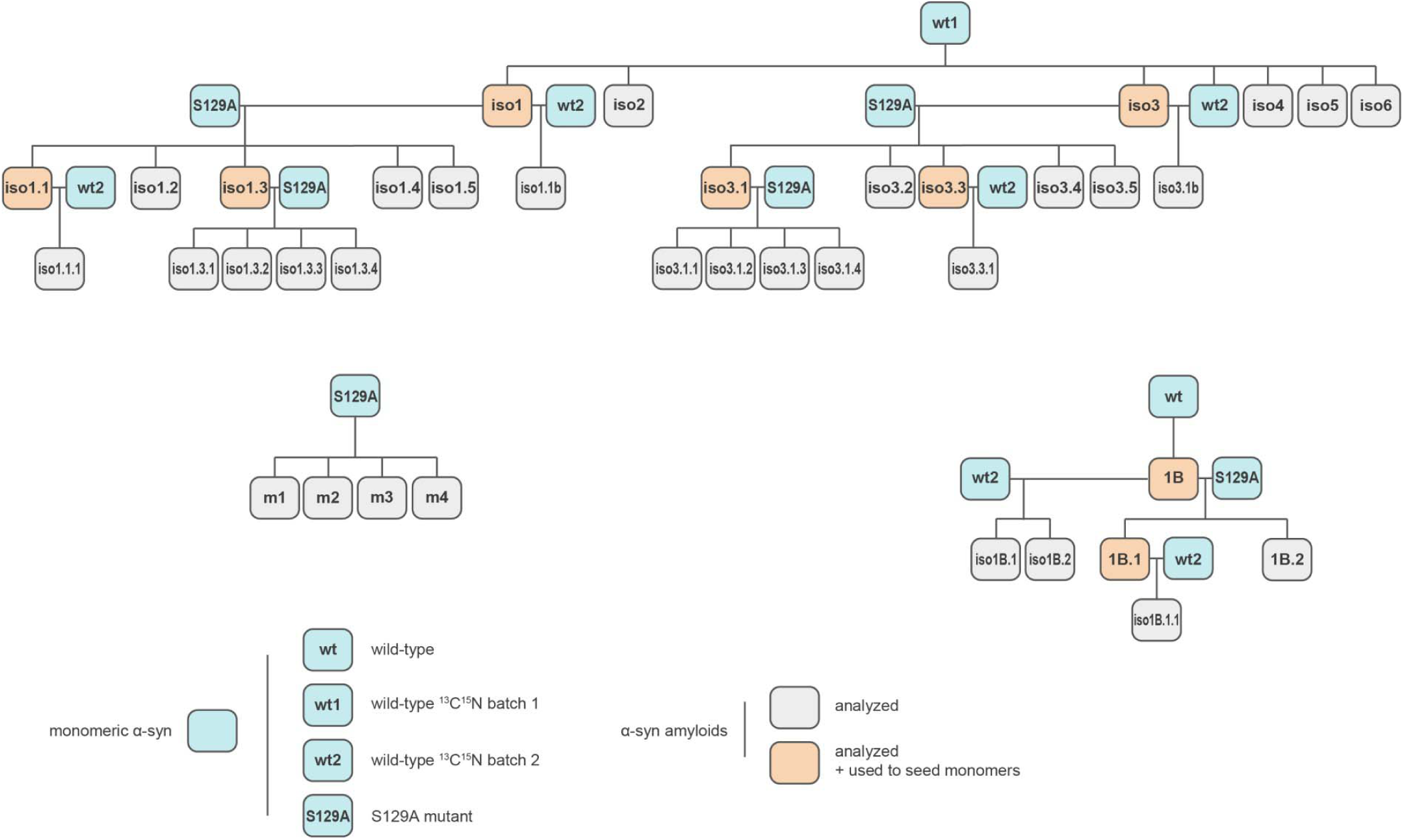
Complete genealogy of the α-syn amyloid polymorphs used in this study.

**Fig. S4.**
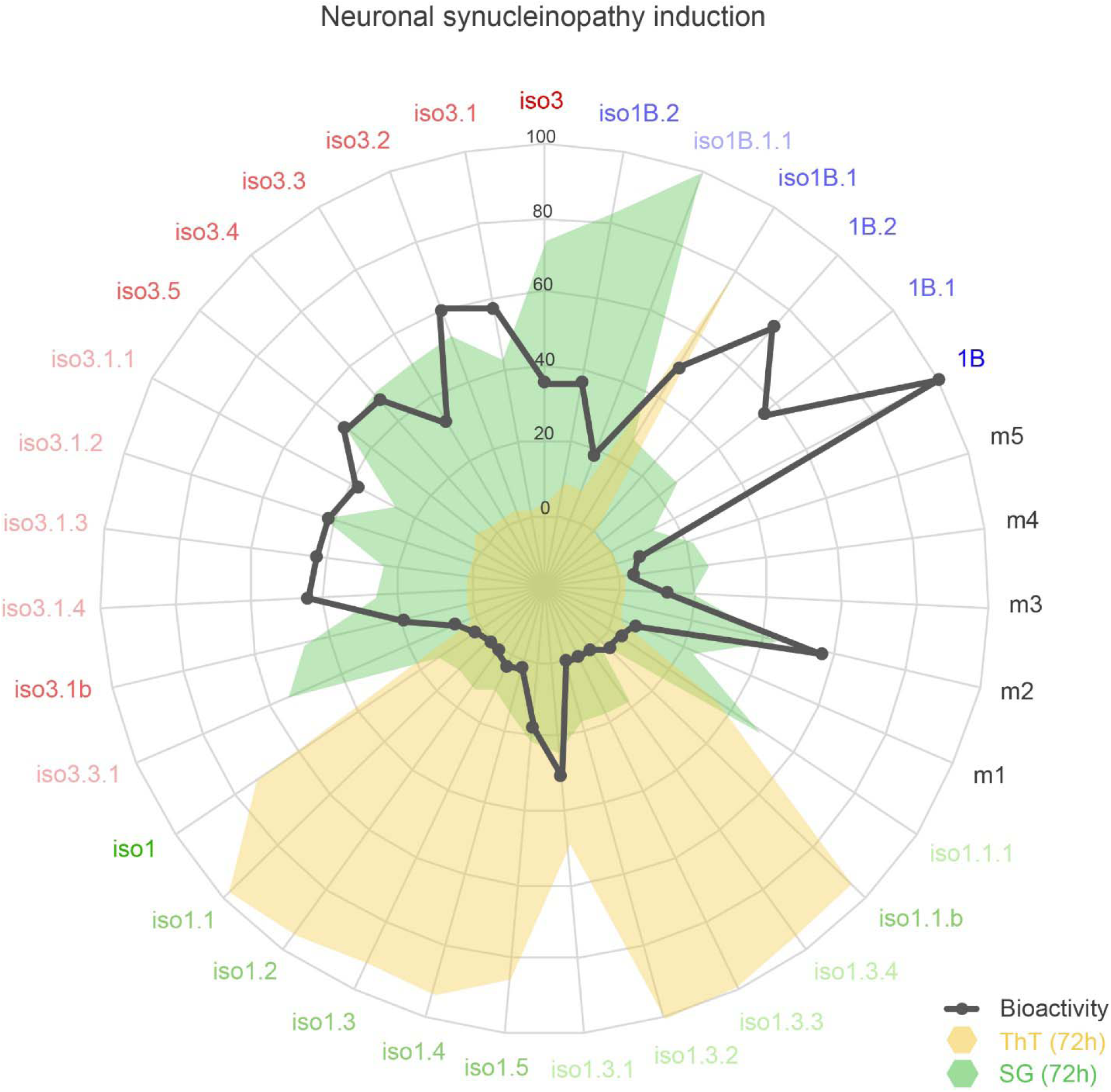
Radar plot of the ThT, SG and neuronal bioactivity values. The members of the genealogy of Fig. S3 and the spontaneous fibrillizations m1 to m5 are shown. For each sample, the three variables were divided by the X-34 value of the sample to normalize the data with respect to the total amyloid contents of the sample.

**Fig. S5.**
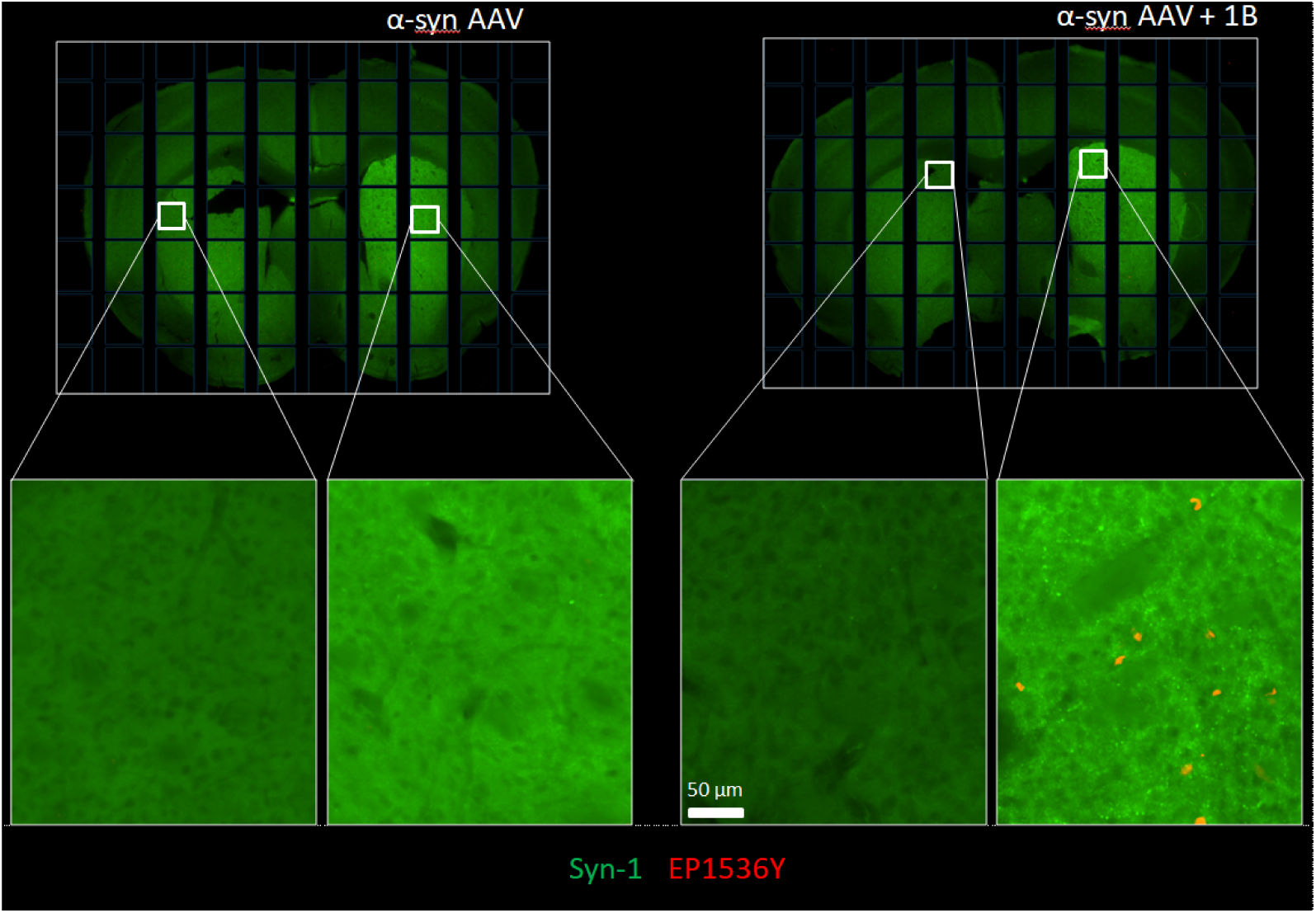
The stealth polymorph 1B triggers a synucleinopathy that spreads along the nigro-striatal pathway *in vivo*. Mice were stereotaxically injected at the level of their right subtantia nigra (SN) with either a bolus of α-syn AVV alone (Left panels), or a bolus of α-syn AAV mixed with stealth polymorph fibrils 1B (Right panels). After four months, immunofluorescence reveals the selective presence of EP1536Y positive α-syn aggregates in the soma of interneurons located in the medio-dorsal region of the striatum, indicating a long distance synucleinopathic spread from the SN towards the rostral striatal regions innervated by the SN neurons.

**Fig. S6.**
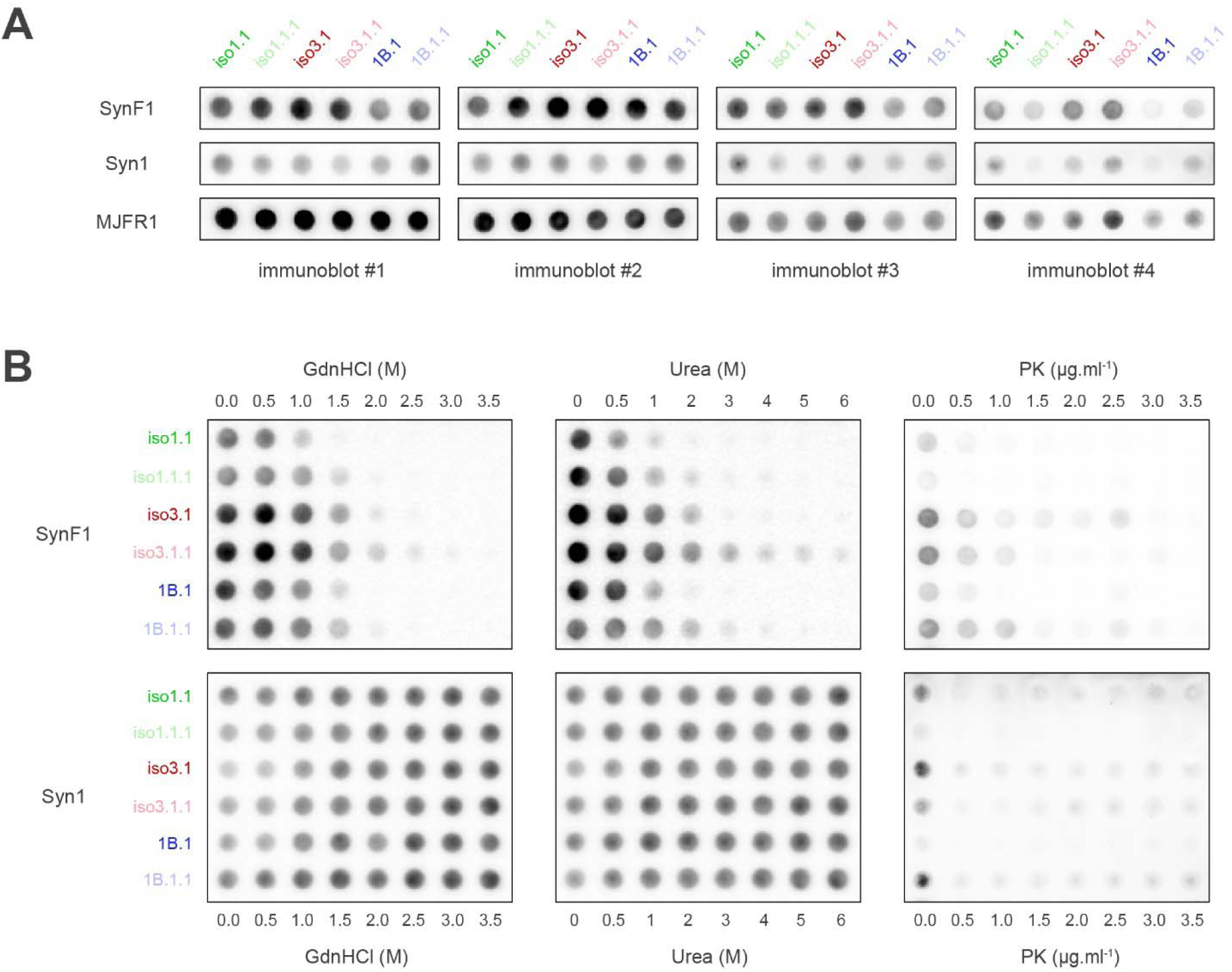
Representative immunoblots of filter trap assays used for the assessment of the fibrils’ immunoreactivity and resistance to disassembly, denaturation and proteolysis. (A) Untouched iso1.1 (dark green), iso1.1.1 (light green), iso3.1 (dark red), iso3.1.1 (light red), 1B.1 (dark blue) and 1B.1.1 (light blue) fibrillar recombinant human α-synuclein preparations were subjected to filter trap assays followed by immunoblotting with antibodies against aggregated (SynF1), monomeric (Syn1) and total human (MJFR1) α-syn. The signal intensities and their ratios were used for quantifying the immunoreactivity of each type of assembly to SynF1. (B) The same fibrillar preparations were treated with indicated increasing concentrations of GdnHCl (1 hour at room temperature), Urea (6 hours at room temperature), and Proteinase K (30 minutes at 37°C). Each sample was subjected to filter trap assay immunoblotted with SynF1 and Syn1 antibodies in order to quantify the disappearance of fibrillar, and the appearance of monomeric α-synuclein forms respectively. The signals normalized to untreated samples allowed the assessment of the resistance of each fibril type to disassembly, denaturation and proteolysis.

**Table S1.** List of the Antibodies used in this study

| Antibody | Target | Company | Cat.No | Dilution IF | Dilution IB |
| --- | --- | --- | --- | --- | --- |
| **Primary antibodies** |  |  |  |  |  |
| MJFR-1 | human alpha-synuclein | Abcam | ab138501 | 1 : 1 000 | 1 : 10 000 |
| EP1536Y | pS129 phospho-synuclein | Abcam | ab51253 | 1 : 500 | 1 : 5 000 |
| Syn1 clone 42 | monomeric alpha-synuclein | BD Biosciences | 610787 | 1 : 500 | 1 : 2 000 |
| Syn-F1 | aggregated alpha-synuclein | BioLegend | 847802 | 1 : 500 | 1 : 10 000 |
| Actin | Beta-actin | Sigma | A5316 | n/a | 1 : 10 000 |
| **Secondary antibodies** |  |  |  |  |  |
| Goat anti-mouse HRP | Mouse IgG (H+L) | Jackson Immuno Research | 115-035-146 | n/a | 1 : 10 000 |
| Goat anti-rabbit HRP | Rabbit IgG (H+L) | Jackson Immuno Research | 111-035-144 | n/a | 1 : 10 000 |
| Donkey anti-mouse Alexa 488 | Mouse IgG (H+L) | Thermo Fisher Scientific | A-21202 | 1 : 500 | n/a |
| Goat anti-rabbit Alexa 488 | Rabbit IgG (H+L) | Thermo Fisher Scientific | A-11008 | 1 : 500 | n/a |
| Donkey anti-mouse Alexa 594 | Mouse IgG (H+L) | Thermo Fisher Scientific | A-21203 | 1 : 500 | n/a |
| Donkey anti-rabbit Alexa 594 | Rabbit IgG (H+L) | Thermo Fisher Scientific | A-21207 | 1 : 500 | n/a |

